# Impact of ultrasonography on identifying noninvasive prenatal screening false-negative aneuploidy

**DOI:** 10.1101/748269

**Authors:** Wei Li, Fanwei Zeng, Baitong Fan, Nan Yu, Jing Wu, Yun Yang, Hui Huang, Sheng-li Li, Zhiyu Peng

## Abstract

**Purpose:** To evaluate the impact of ultrasonography (US) on identifying noninvasive prenatal screening (NIPS) false-negative aneuploidy.

**Methods:** Analysis of large population-based NIPS false-negative aneuploidy data comprising karyotypes, clinical outcomes, and US results.

**Results:** From December 2010 to July 2018, a total of 3,320,457 pregnancies were screened by NIPS performed in BGI; among them, 69 NIPS false-negative aneuploidy cases with informed consent were confirmed, and US examination data for 48 cases were not available. Of the 21 cases with US results, 19 (90.5%) had various abnormalities on ultrasound, and 2 (9.5%) cases were shown to be normal on ultrasound. Additionally, 6 out of 7 live born fetuses (approximately 85.7%) were found to have abnormalities on ultrasound. Ventricular septal defects constituted the most frequently observed ultrasound abnormality type among the 21 NIPS false-negative aneuploidy cases.

**Conclusion:** NIPS has expanded rapidly worldwide and now accounts for a large proportion of prenatal screening tests in China. This study suggests that abnormal US findings should not be neglected, even when NIPS produces a negative result. Combining NIPS with an US examination can further reduce the incidence of livebirths with aneuploidy.

## INTRODUCTION

Noninvasive prenatal screening (NIPS) has expanded rapidly worldwide and now accounts for a large proportion of prenatal screening tests for chromosomal malformations.^1–3^ By means of cell-free DNA genomic sequencing analysis, NIPS for trisomies 21, 18, and 13 achieves much better performance than conventional standard screening tests, which are based on serological markers, maternal age, and maternal history.^3^ The American College of Medical Genetics and Genomics has recommended replacing traditional biochemical screening tests with NIPS for trisomies 21, 18, and 13 across the maternal age spectrum.^4^ The potential impact of NIPS on the field of prenatal diagnosis and on the prevalence of livebirths with chromosomal abnormalities is increasing dramatically since sequencing costs have been gradually decreasing and since government funding has increased.^5^

Although the occurring rate of false-negative cases is very low, discordant findings between NIPS and prenatal diagnoses, which may result from a low fetal DNA fraction or fetoplacental mosaicism,^6,7^ are still observed globally. There are case reports that ultrasonography (US), a powerful method for screening fetal aneuploidy during pregnancy, may contribute to the identification of NIPS false-negative cases.^8,9^ Abnormalities on US are useful markers for detecting trisomy 21, trisomy 18, trisomy 13, sex chromosome aneuploidy (SCA), rare autosomal trisomies (RATs) and even copy number variations, as reported previously.^10–12^ However, reports on the quantitative contribution of US toward identifying NIPS false-negative cases are still limited.

The NIPS result represents the genotype of the fetus, whereas the US finding reflects the phenotype of the fetus. We aimed to assess the impact of US on noninvasive prenatal screening false-negative aneuploidy based on real-world data from 3,320,457 pregnancies screened from 2010 to 2018.

## MATERIALS AND METHODS

### Study population and sample collection

From December 2010 to July 2018, a total of 3,320,457 pregnancies were screened by NIPS performed by BGI, among which 69 NIPS false-negative aneuploidy cases with informed consent for research purposes were confirmed. This study was approved by the institutional review board of BGI (BGI-IRB).

NIPS and US examination were recommended for pregnancies according to the standard screening process^13^ ; thereafter, G-banded karyotyping or a chromosomal microarray (CMA) diagnosis was highly recommended for cases that were high risk.

### Validation and follow-up of NIPS false-negative aneuploidy cases

The reporting of false-negative NIPS results (for trisomy 21, 18, 13) was encouraged by offering each participant insurance as part of the test. For each false-negative NIPS case with clinically confirmed aneuploidy by G-banded karyotyping or CMA diagnosis before or after a live birth, the insurance policy requires a payment of CNY 20,000 or CNY 400,000, respectively, to the patient. The ultrasound examination results, if available, could be traced from the insurance materials. The unavailability of ultrasound information meant either that US was not performed or that the US results were not available from the insurance materials.

#### Statistical analyses

Statistical analysis was performed using Student’s *t*-test, and *P* < 0.05 was considered statistically significant. All statistical analyses were conducted using R, version 3.4.3.

## RESULTS

The enrollment, clinical follow-up and outcomes of the false-negative cases participating in NIPS and US are presented in **Figure 1**. Of the 69 NIPS false-negative pregnancies (**Table S1**), 21 cases had available ultrasound examination information, and 48 cases did not have available information. The median gestational age (GA) of the 21 NIPS false-negative cases that underwent US examinations was 24.8 weeks; herein, the second-trimester gestational age group (66.7%) was the most predominant (**Table S2**). The median maternal age was 30 years, and most of the women, 12 out of 21 cases (57.1%), were not more than 30 years old (**Table S2**).

**Figure 1.**
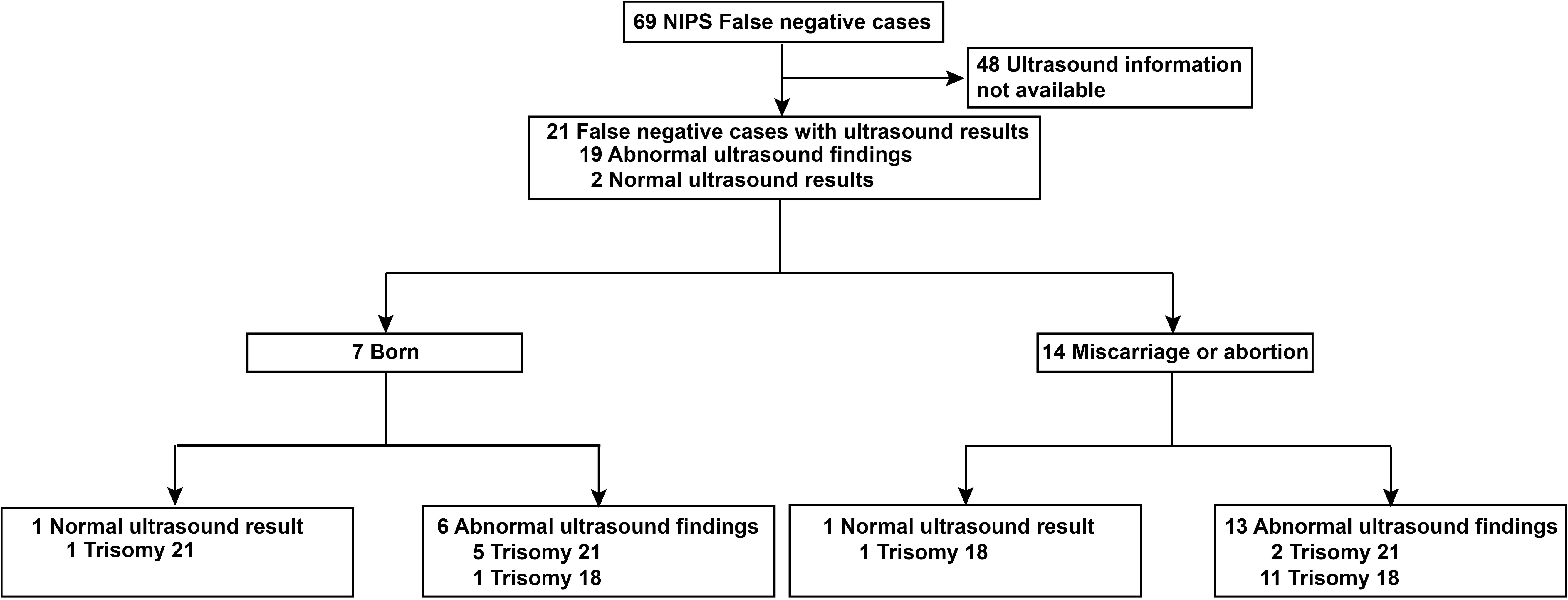
Enrollment, clinical follow-up and outcome classification of the false-negative cases participating in the NIPS and US examinations.

Regarding the 21 cases with US information (**Figure 1**, **Table 1**), a total of 19 pregnancies were found to have abnormalities by US examination, whereas only 2 pregnancies were normal by ultrasound examination (**Figure 1**). That is, 90.5% of the cases with ultrasound information (19 out of 21) and at least 27.5% of the total cohort (19 out of 69) were NIPS false-negative cases that could potentially be avoided by means of a combination of NIPS and an US examination, followed by G-banded karyotyping or CMA diagnosis, which would probably not increase the risk of fetal loss.^14^ It did not escape our notice that 6 out of 7 born cases had abnormal ultrasound findings during pregnancy; that is, 6 out of 7 born NIPS false-negative cases could have potentially been identified by means of the US examination (**Figure 1, Table 1**). The abnormal ultrasound findings for the 6 born cases included small head circumference, lateral ventriculomegaly (11.2/10.2 mm), ventricular septal defect, aortarctia, tricuspid moderate regurgitation, nasal bones dysplasia, short femora, short humeri, bilateral renal pelvis separation (4.9/4.0 mm), polyhydramnios, and single umbilical artery (**Table 1**).

The features of abnormal ultrasound findings and the corresponding karyotyping genotypes for the 19 cases (**Table 1**) were analyzed and summarized in **Figure S1**. Congenital cardiovascular defects, skeletal malformations, and brain and nervous system defects were the most frequently detected organic system abnormalities (**Figure S1a**), and ventricular septal defects (7 out of 19 cases) were the most frequently observed ultrasound abnormalities (**Figure S1a**) among the NIPS false-negative aneuploidy cases. There is a preponderance of the trisomy 18 genotype over the trisomy 21 genotype among all of the top six ultrasound-based systematic abnormalities, including the cardiovascular, skeletal, brain, and nervous, urinary, fetal appendage, and craniofacial systems (**Figure S1a**). Among the defects in the cardiovascular system, ventricular septal defect, pulmonary artery enlargement, dilated right heart, hypoplastic left heart, and double outlets of the right ventricle could be strong ultrasonic markers for trisomy 18, whereas an atrioventricular septal defect suggests trisomy 21. Additionally, tricuspid regurgitation and aortarctia could be ultrasonic findings for either trisomy 18 or trisomy 21. As shown in **Figure S1b**, the fetuses with trisomy 18 tended to show abnormal findings by ultrasound at an earlier average GA (24.5 versus 27.8 weeks) than those with trisomy 21; however, the difference was not significant (*P* value was 0.19).

## DISCUSSION

With the largest population-based data to date, this study analyzes the impact of prenatal US examination on NIPS false-negative cases. This report focuses on cases between December 2010 and July 2018 in China, and these data may also have great significance in other countries as a reference. Even though the occurring rate of NIPS false-negative cases was low, we aimed to investigate the false-negative case data and explore a viable method to supplement NIPS to further reduce the livebirth prevalence of aneuploidy.

### Combination NIPS and US

NIPS is the most accurate and powerful prenatal screening method for Patau, Edwards, and Down syndromes to date, according to a previous global report.^4,15^ Nevertheless, NIPS can still produce false-negative chromosomal aneuploidy results for fetuses, as well as livebirths when the probability of a livebirth (approximately 20% of trisomy 21 fetuses may progress to term delivery)^16^ is taken into account. NIPS false-negative findings may occur due to various reasons, including a low fetal DNA fraction and fetoplacental mosaicism. As was reported previously, mosaic trisomy can be accurately detected by NIPS only when the fraction of fetoplacental mosaicism is higher than 70%.^6^

The combination of NIPS and an US examination may further reduce the risk for false-negative fetuses and livebirths. We observed a significant proportion (90.5%, 19 out of 21 cases with ultrasound data) of NIPS false-negative cases (at least 27.5% of the total cohort [19 out of 69] if all of the US cases without available data had normal US results or 97.1% [67 out of 69] if all of the US cases without available data had abnormal US results) could be potentially avoided by means of a combination of NIPS and an US examination, followed by G-banded karyotyping or CMA diagnosis, which would probably not increase the risk of fetal loss, according to an updated large population study.^14^ Clinicians should pay additional attention to the top-ranked US findings indicating congenital cardiovascular defects, skeletal malformations, brain and nervous system defects, urinary malformations, and especially ventricular septal defects, which are the most common abnormal findings on US among NIPS false-negative cases (**Figure S2**). In addition, special attention should be paid to abnormal signs on US, namely, small head circumference, lateral ventriculomegaly (11.2/10.2 mm), ventricular septal defect, aortarctia, tricuspid moderate regurgitation, nasal bones dysplasia, short femora, short humeri, bilateral renal pelvis separation (4.9/4.0 mm), polyhydramnios, and single umbilical artery, since six NIPS false-negative fetuses (five trisomy 21 cases and one trisomy 18 case) with these US signs were born alive because these abnormal US findings were overlooked (**Table 1**). Five of these cases (cases 2, 3, 4, 5, and 7) with strong ultrasound markers (in isolation or with additional anomalies) would be advised by clinicians to undergo an invasive diagnosis,^17^ while the other case (case 1) with a soft ultrasound marker may not be advised to undergo an invasive diagnosis by the clinician in China. Reasonably, all five fetuses with soft markers (ultrasound findings), including small head circumference, nasal bone dysplasia, lateral ventriculomegaly (11.2/10.2 mm), polyhydramnios, and bilateral renal pelvis separation (4.9/4.0 mm), successfully progressed to term delivery, indicating that these types of ultrasound findings are inclined to be overlooked more frequently (**Table 1**).

This implies the complementary role of ultrasonography with NIPS to achieve a better prenatal screening performance. Clinically, abnormal US findings should not be neglected when performing G-banded karyotyping or the CMA diagnosis, even when NIPS results in a negative result.

Our study had the following limitations. First, the US results were not available for all of the 69 cases, so we could not accurately evaluate the quantitative contributions of US toward identifying NIPS false-negative cases. The rate of abnormal ultrasound findings among the NIPS false-negative cases may be approximately 90.5% (19 out of 21 cases with US data), fluctuating between 27.5%, (at least 19 out of 69, if all of the US cases without available data had US normal results) and 97.1% (at most 67 out of 69, if all of the US cases without available data had US abnormal results). Second, accurate follow-up of some clinical information was not performed. Specifically, the reasons that 6 cases with prenatal abnormal findings on ultrasound were born were undetermined in our study, and these reasons may be that the clinicians overlooked the findings or that the patients were unwilling to undergo abortions. On the latter situation, US may not be helpful for further reducing the livebirth incidence of false-negative aneuploidy cases. Lastly, detailed clinical ultrasound information, such as the presence of multiple malformations (Table 1), was not available for all the cases because these details were not specified in the insurance materials.

## Conclusion

NIPS has expanded rapidly worldwide and now accounts for a large proportion of prenatal screening tests in China. We observed that 19 out of the 69 NIPS false-negative pregnancies were found to have abnormal findings on ultrasound, indicating that between at least 27.5% (if all of the US cases without available data had normal US results) and at most 97.1% (if all of the US cases without available data had abnormal US findings) false-negative cases could be potentially avoided by means of a combination of NIPS and ultrasonography, followed by G-banded karyotyping or a CMA diagnosis. Based on all the information presented above, combining NIPS and ultrasonography can potentially reduce the livebirth incidence of false-negative aneuploidy cases.

## SUPPLEMENTARY INFORMATION

The online version of this article contains supplementary material, which is available to authorized users.

## ACKNOWLEDGMENTS

We thank all medical centers, participants, and the families in this study for their cooperation and support.

## FIGURE LEGENDS

**Figure S1.**
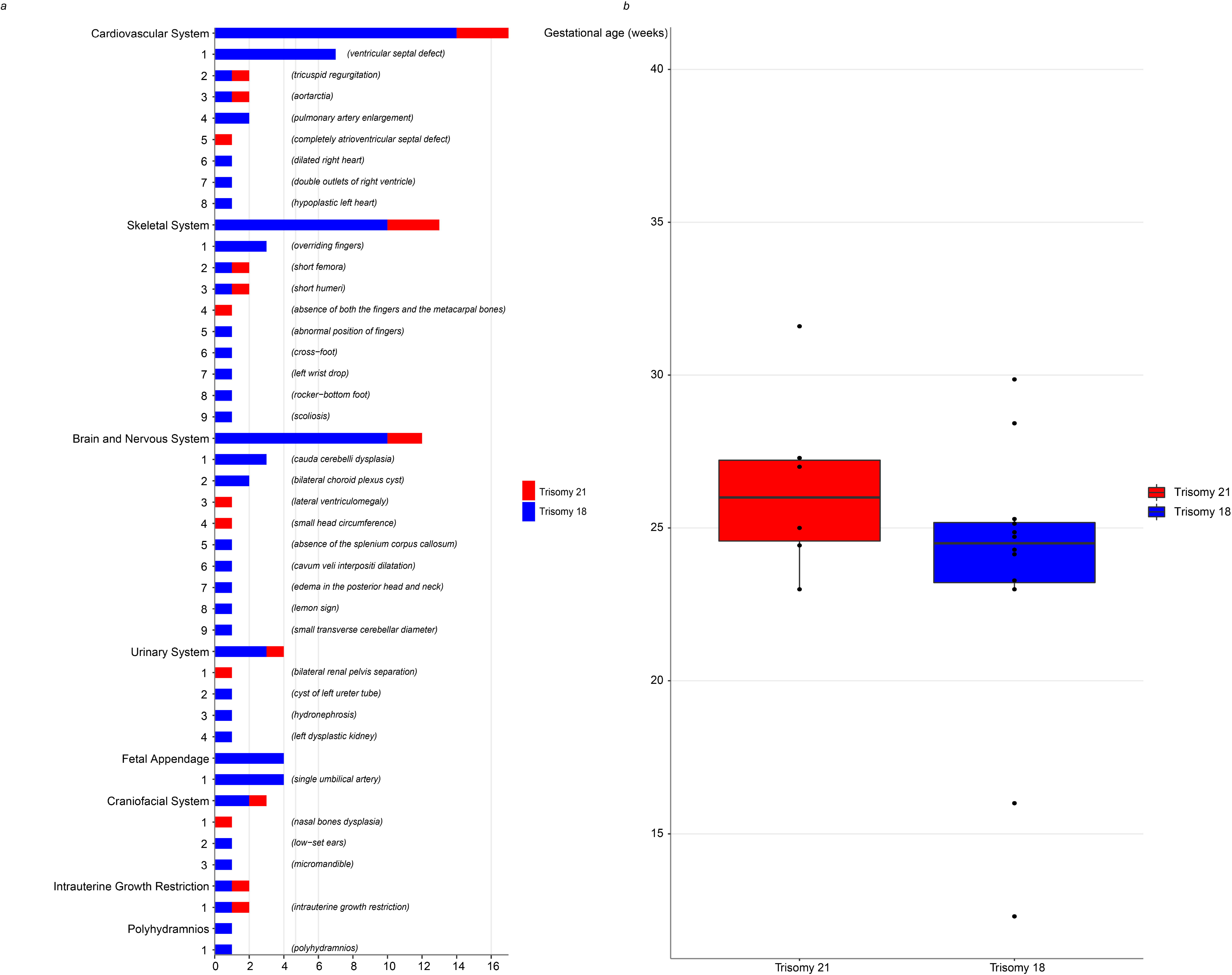
The distribution of abnormal ultrasound phenotypes. (a) Histogram showing the number of distinct ultrasound abnormal phenotypes based on affected organ systems, as well as the distinct ultrasound findings and corresponding genotypes. (b) Boxplot comparison of the gestational age for the abnormal ultrasound findings among 5 trisomy 21 fetuses and 11 trisomy 18 fetuses. Statistical analysis was performed using Student’s t-test, and the *P* value was 0.19.

**Figure S2 Ultrasonography.**

**Case 8 Absence of the fingers and the metacarpal bones of the right hand**

A. The distal right forearm of the fetus was not evident by two-dimensional ultrasound. B. Absence of the right finger bone and metacarpal bone by three-dimensional ultrasound. The fetal condition was complicated by intrauterine growth restriction, and the chromosome karyotype was confirmed to be trisomy 21. R-ARM, right forearm.

**Case 9 Complete atrioventricular septal defect**

The fetus had a complete atrioventricular septal defect, and the chromosome karyotype was confirmed to be mos 47, XN, +21[45]/46, XN [40]. AVSD, atrioventricular septal defect; VS, ventricular septum; AS, atrial septum.

**Case 19 Choroid plexus cyst**

The cyst mass can be seen in the bilateral choroid plexus of the fetus. The fetal condition was complicated by a small transverse cerebellar diameter, cavum veli interpositi dilatation, short femora, short humeri, and intrauterine growth restriction. The chromosome karyotype was confirmed to be trisomy 18. CPC, choroid plexus cyst.

