## Supplemental Table 2 for "Impact of ultrasonography on identifying noninvasive prenatal screening false-negative aneuploidy"

**Table S2 Demographic and Characteristics of the 21 false negative cases with US results**

| **Characteristic** | **Value** |
| --- | --- |
| Median maternal age (IQR) -years | 30.0 (28.0-33.0) |
| ≤ 30 years (%) | 12 (57.1%) |
| 31-34 years (%) | 4 (19.0%) |
| ≥ 35 years (%) | 5 (23.8%) |
| Median gestational age of abnormal US findings (IQR) -weeks | 24.8 (23.5-26.6) |
| First trimester (9-13 weeks) (%) | 1 (4.8%) |
| Second trimester (14-27 weeks) (%) | 14 (66.7%) |
| Third trimester (≥ 28 weeks) (%) | 3 (14.3%) |
| Information not available (%) | 3 (14.3%) |
| Fetuses genotypes |  |
| Trisomy 21 (%) | 8 (38.1%) |
| Trisomy 18 (%) | 13 (61.9%) |
| Trisomy 13 (%) | 0 (0%) |
| Mosaicism (%) | 1 (4.8%) |
| Fetuses clinical outcome |  |
| Abortion | 13 (61.9%) |
| Miscarriage | 1 (4.8%) |
| Born | 7 (33.3%) |
| US findings |  |
| Abnormal (%) | 19 (90.5%) |
| Normal (%) | 2 (9.5%) |

IQR, interquartile range.
